# Photorelease of Diacylglycerol Increases the Amplitude and Duration of Protein Kinase C-BetaII Relocation *in cyto*

**DOI:** 10.1101/102657

**Authors:** Joachim Goedhart, Theodorus W.J. Gadella

## Abstract

Diacylglycerol (DAG) is a lipid second messenger produced by receptor stimulated phospholipase C and is capable of activating several PKC isoforms. Classical PKC isoforms require simultaneous presence of calcium and DAG for activation and relocation to membranes. The aim of this study was to synthesize a photolabile precursor of DAG and examine the effect of an immediate increase of the signaling lipid on PKC relocation. Caged DAG was synthesized using a photoreleasable 7-diethyl-aminocoumarin (DEACM) group. Photolysis was monitored *in vitro* by an increase in coumarin fluorescence from which an uncaging quantum yield of 1.1% was determined. This quantum yield proved ideal for efficient uncaging at high UV power while allowing localization of the fluorescent compound at low UV power. Taking advantage of the coumarin fluorescence, it was demonstrated that DEACM-DiC8 accumulated inside cells. Effects of DAG photorelease on periodic relocations of PKCbetaII, induced by histamine, UTP or EGF, were studied. Photorelease of DAG *in cyto* immediately increased the amplitude and duration of relocation events, regardless of the agonist used. Together, the results demonstrate the usefulness of caged DAG for dissecting PKC signaling and suggest that DAG levels are limiting during signaling.

## INTRODUCTION

Diacylglycerol (DAG) is a lipid second messenger of which the levels can be increased during cellular signaling. The classical pathways that produce DAG involve the activation of a phosphoinositide specific phospholipase C (PLC), either by activation of the gamma isoforms by receptor tyrosine kinases (RTK) or by activation of the beta isoforms by G-protein coupled receptors (GPCR) [1]. In both pathways, the activated PLC hydrolyses phosphatidylinositol-4,5-bisphosphate yielding DAG and the soluble second messenger Ins(1,4,5)P3, of which the latter triggers an elevation of the intracellular calcium concentration [2].

Simultaneous presence of calcium and DAG is required for the activation of classical PKC isoforms [3]. Experiments using GFP-tagged PKCs have addressed the spatiotemporal regulation of classical PKC relocation and activation in single living cells [4-6]. These studies have led to a model for PKC activation in which several steps can be discerned. The relocation of PKC is initiated by association of the calcium-sensitive C2-domain with the plasma membrane upon elevation of calcium levels. Next, a conformational change allows binding of the C1-domains to DAG. Binding of both the C1-domain to DAG and the C2-domain to calcium is necessary to release the pseudosubstrate from the active site [3]. Upon release of the pseudo-substrate the active site is exposed and the enzyme can exert its kinase activity on substrates.

To understand the regulation of PKCs by calcium and DAG, several approaches are used. An elegant method to achieve control over the spatial and temporal concentration of second messenger is to use caging technology [7]. The part of the molecule that is necessary for its bioactivity is masked by a protecting group. This protecting group can be removed by irradiation with low-energy UV light (around 360 nm). Caged calcium, or caged calcium scavengers are commercially available and have been successfully applied in numerous studies. These tools were used to examine the role of calcium in controlling PKC location and activity in single living cells [8, 9]. A similar approach using caged analogs of DAG to study the effect of DAG on PKC relocation is still lacking. Previously, caged dioctanoylglycerol (DiC8) has been used by Walker and colleagues to study the effect of DAG on peak calcium current and on the regulation of the twitch amplitude in single cardiac myocytes [10-12]. Although there is compelling circumstantial evidence that the observed effects of DAG are mediated by PKC, direct effects on PKC, i.e. relocation or activation, were not studied.

In order to be able to assess the role of DAG in regulating the activity and location of a classical PKC in single living cells, we synthesized and characterized a caged DAG analog. The caging moiety is a coumarin-derivative [13], which has a relative high extinction coefficient and can absorb light of wavelengths >400 nm. Another advantage is the intrinsic fluorescence, which in some cases is increased upon uncaging, allowing easy analysis of the uncaging reaction [14]. The photoreleasable DAG was used to evaluate the role of DAG on the relocation of PKCbetaII during signaling.

## MATERIALS AND METHODS

### Synthesis

7-diethylamino-4-hydroxymethylcoumarin (DEACM-OH) [15] was activated with 4- nitrophenylchlorformate as described [16]. This compound was reacted with solketal from which the caged diacylglycerol was synthesized as described [17].

### Spectroscopy and Quantum yield

The photon flux at 360 nm (slit 10 nm) was determined by ferrioxalate actinometry as detailed before [18]. Identical settings were used to measure the fluorescence increase of DEACM-DiC8 using a PTI fluorimeter. The liberation of the cage was assayed by monitoring fluorescence at 450 nm (slit 2 nm) while stirring. The concentration of DEACM-DiC8 was 0.23 *µ*M to meet the requirement that εcl<0.01 allowing determination of the quantum yield independent of the concentration [18]. Experiments in micelles were performed using 0.2 % (v/v) Triton X-100 and 20 mM Hepes (pH=7.4)

### Cell culturing and transfection

HeLa cells were seeded in 6-well plates containing 25 mm Ø coverslips, thickness #1. The next day, cells were transfected with 1 *µ*g of plasmid DNA encoding C1a-EGFP (a kind gift of Tobias Meyer, Stanford University, Stanford, CA) or EGFP-PKCbetaII (a kind gift of Yusuf Hannun, Duke University Medical Center, Durham, NC). One day after transfection the coverslip was mounted into a home made chamber and submerged in 1 ml of extracellular medium (20 mM HEPES (pH=7.4), 137 mM NaCl, 5.4 mM KCl, 1.8 mM CaCl2, 0.8 mM MgCl2 and 20 mM glucose).

### Fluorescence microscopy

HeLa cells were observed and imaged on a LSM510 (Zeiss) confocal laser scanning microscope. A Zeiss 63× water-immersion objective (C-Apochromat, NA 1.2) with correction ring was used for scanning samples. EGFP fluorescence was excited using the 488 nm argon ion laser line (AOTF maximally at 5%). The resulting fluorescence was separated from the excitation light with a 488 nm dichroic mirror and passed through a longpass filter (LP505 nm). The pinhole was set corresponding to 1-2 airy units to obtain confocal sections. Coumarin was excited using the 351/364 nm laser lines of a Innova Enterprise II ion laser (Coherent, Santa Clara, CA), which was reflected onto the sample by a multiband dichroic mirror (UV/488/543/633 nm), allowing simultaneous excitation of EGFP at 488 nm. Coumarin fluorescence emission was filtered through a 385 nm longpass or a 380-430 nm bandpass filter. For uncaging the sample was exposed to a single scan (<1s) with the 351/364 nm laser (AOTF at 100%).

## RESULTS

### Synthesis and spectral properties of DEACM-DiC8

Previously, the synthesis of a related coumarin-caged DAG has been described [12, 19]. Our synthesis approach is different in that we react the caged group to solketal rather than direct coupling to DAG. After deprotection, this allows a variety of fatty acids to be attached to the glycerol moiety. In addition, the product is not contaminated by unreacted DAG. This approach is similar to the synthesis of a nitrophenyl caged DAG [17]. The structure of the resulting compound, 7-diethylaminocoumarin caged 1,2-dioctanoyl-*sn*-glycerol (DEACM-DiC8) is shown in figure 1A.

**Figure 1.**
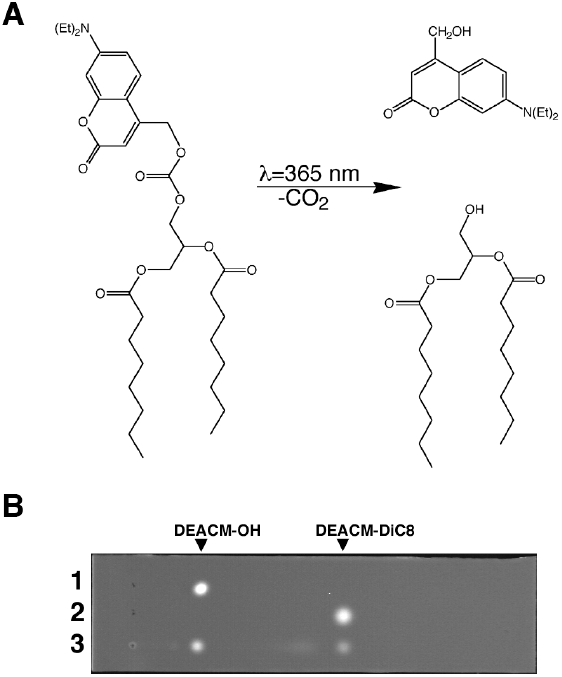
(A) Structure of DEACM-DiC8 and the products, DEACM-OH and 1,2-dioctanoyl-*sn*-glycerol, after photolysis. (B) TLC analysis of the uncaging reaction. DEACM-DiC8 (around 2 nmol) was spotted on the TLC in lane 3 and subsequently exposed to 365 nm light on a transilluminator UV tray for 1 min. Then DEACM-DiC8 (lane 2) and the reference compound DEACM-OH (lane 1) were spotted and the compounds were separated in ethylacetate-hexane (1:1). Compounds were visualized by capturing their fluorescence on a UV tray (312 nm excitation)

Due the intrinsic fluorescence of 7-diethylaminocoumarin, the caged compound can easily be detected on TLC (fig. 1B). After exposing DEACM-DiC8 to UV light (365 nm) the mobility of the resulting compound on TLC is markedly reduced and corresponds to 7-diethylamino-4-hydroxymethylcoumarin (DEACM-OH), indicating that the coumarin cage can be released from DEACM-DiC8 upon UV absorption.

### Spectroscopic characterization of DEACM-DiC8

Coumarin cages in general and the DEACM cage in particular is known for its absorbance at high wavelengths [15, 20]. This is also apparent from the absorbance spectrum of DEACM-DiC8 (fig. 2), showing a typical coumarin shape with a maximum at 380 nm. The coumarin still has a significant absorbance around 400 nm, which allows photolysis of this compound with visible light. The fluorescence emission is maximal at 476 nm.

**Figure 2.**
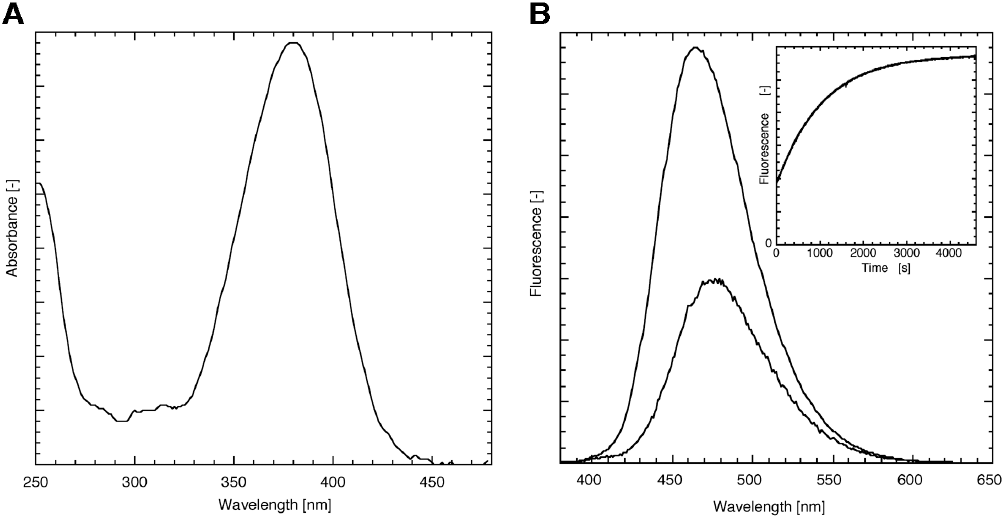
Spectroscopic characterization of DEACM-DiC8 in EtOH. (A) The absorbance of the caging moiety shows a typical coumarin shape. (B) Fluorescence emission spectrum of the DEACM-DiC8 before (λ_max_ = 476 nm) and after photolysis (λ_max_ = 464 nm) acquired at an excitation wavelength of 360 nm show that the fluorescence intensity is increased upon photorelease. The inset of panel B shows the time-dependent increase in fluorescence (excitation at 360 nm, emission at 450 nm), which can be fit to a mono-exponential increase with a rate of 0.95×10^-3^ s^-1^ (R>0.999).

Several coumarin derivatives are known to increase their fluorescence upon photolysis [21, 22]. This is a useful property to directly monitor and quantify photolysis [14]. So far, a similar fluorescence increase for DEACM caged compounds was not reported. Ethanolic solutions of DEACM-DiC8, illuminated on a UV tray with 365 nm light showed an increase in fluorescence. Besides the fluorescence increase after UV exposure, the fluorescence emission maximum of the sample is shifted to 464 nm (figure 2). To study the kinetics of the fluorescence increase, the fluorescence of DEACM-DiC8 was acquired using a fluorimeter by continuous excitation at 360 nm and monitoring the fluorescence at 450 nm. The fluorescence increase in EtOH was determined to be at least 3-fold (inset of fig. 2B): 3.3 ± 0.2 (n=3). When the experiment is performed in Triton X-100 micelles, mimicking an environment with a water-membrane interface, similar curves were obtained with only a modest 1.4-fold increase.

Previously, we have reported an assay for directly monitoring the photorelease of a caged phosphatidic acid. The equations describing the fluorescence increase that accompanied the photorelease were based on a compound with two caging moieties [18]. The caged DAG has only one cage and the equation describing the release of free DEACM and DiC8 is simplified to a mono-exponential increase of compound during illumination. By curve fitting the fluorescence increase a rate of 0.95×10^-3^ s^-1^, was obtained corresponding to a half-time of 730 s. To determine the uncaging quantum yield, the photon flux was measured at 360 nm by ferrioxalate actinometry under exactly the same circumstances as the DEACM-DiC8 uncaging experiment, which gave 4.3 10^15^ s^-1^. From the uncaging rate, the photon flux and the extinction coefficient the quantum yield can be calculated as described before [18], yielding a value of 1.1 %. The reported quantum yield of a related coumarin-DAG varies from 1.4% [19] to 40% [12]. The quantum yield determined in this study is close to the value reported by Suzuki et al., but greatly differs from the value determined by Robu et al. for unknown reasons. The quantum yield of 1.1% is high enough for efficient uncaging, yet it is low enough for enabling routine observation of the coumarin fluorescence without triggering too much DAG production. In this respect it is of note that a quantum yield of 40% would not be compatible with observation of fluorescence.

### Detection of DAG photorelease in vivo

The TLC analysis shown in fig. 1B demonstrates the photolysis of DEACM-DiC8 in vitro and the release of the fluorescent protecting coumarin. However, the formation of DiC8 cannot be detected on this TLC. Since it has been shown that a fusion protein comprising the first C1-domain of PKC-γ and Enhanced Green Fluorescent Protein (C1a-EGFP) can be used as a sensor for DAG in single living cells [23], we wanted to employ this system to detect the photoreleased DAG *in cyto*.

First, the detection sensitivity of the DAG analog DiC8 was characterized. HeLa cells were exposed to different concentrations of DiC8 and after 5 minutes the distribution of C1a-EGFP in living cells was acquired by confocal laser scanning microscopy. As shown in figure 3, cells that are not incubated with DiC8 show homogeneous distribution of C1a-GFP throughout the cytosol and nucleus. In cells that are incubated with 10 *µ*M DiC8 a clear membrane localization of C1a-EGFP was observed with a concomitant depletion of fluorescence in the cytosol. Even at a 10-fold lower concentration, still an increase of fluorescence at the plasma membrane relative to the cytosol can be detected.

**Figure 3.**
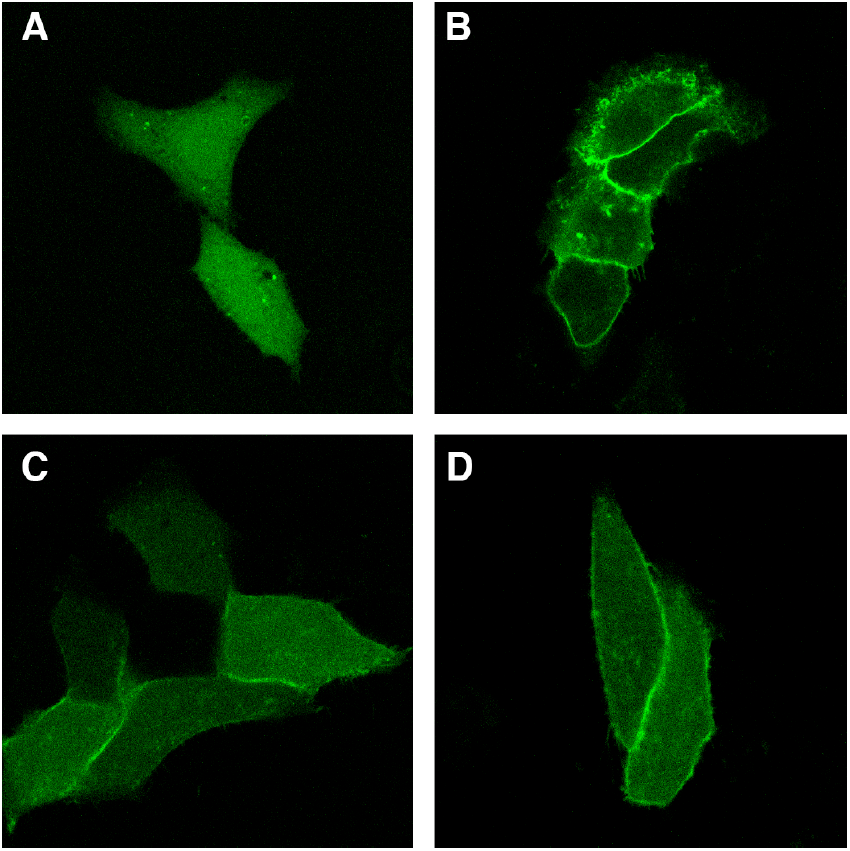
Location of C1a-GFP in living HeLa cells and relocation by addition of DiC8. Hela cells expressing C1a-GFP show homogeneous distribution of the fluorescence in the cytosol and nucleus (A). Upon addition of 10 *µ*M DiC8 (B), the fluorescence is increased at the plasma membrane and almost completely depleted from the cytosol. At a lower concentrations (1 *µ*M) of externally added DiC8, the relocation is reduced, but still clearly visible as accumulated fluorescence at the plasma membrane (C). When 6 *µ*M photolysed DEAM-DiC8 is added to cells expressing C1a-GFP, the fluorescence is increased at the plasma membrane, indicating the release of DAG from its caged precursor (D).

Next, we incubated cells with 6 *µ*M DEACM-DiC8 that was either added directly, or, exposed to UV to prior to adding it to cells. Cells treated with non-exposed caged compound did not show a significant membrane accumulation of the DAG sensor (data not shown). In contrast, when cells were incubated with photolysed DEACM-DiC8 a clear accumulation of C1a-GFP at the plasma membrane was observed, as shown in figure 3D. This experiment shows that during UV exposure DiC8 is released from DEACM-DiC8 and that it can be detected by living cells, whereas the intact caged compound is not detected by cells.

### Incorporation of DEACM-DiC8 in cells

To examine whether DEACM-DiC8 is incorporated into cells, cells were incubated with 200 *µ*M caged DAG and visualized by fluorescence microscopy. From figure 4A it can be inferred, that the probe is located in the cytosol and excluded from the nucleus. The reticular distribution of the cytosolic fluorescence suggested that the probe is predominantly associated with endomembranes. Besides a relative equal distribution of fluorescence throughout the cells we consistently observed intense blue fluorescent spots. To examine whether these structures correspond to lipid droplets, a dual labeling with Nile red was performed [24]. As can be inferred from fig. 4A and 4B, there is a strong colocalization of the Nile red and coumarin fluorescence, indicating that DEACM-DiC8 accumulates in lipid droplets. Strikingly, when cells were illuminated with UV light for photolysis of DEACM-DiC8, the highly fluorescent spots disappeared (fig. 4C and 4D). We conclude that the DAG moiety is responsible for the location of the caged compound in lipid droplets. Apparently, the hydrophilic caging moiety is released from the lipid droplets once it is cleaved from DAG.

**Figure 4.**
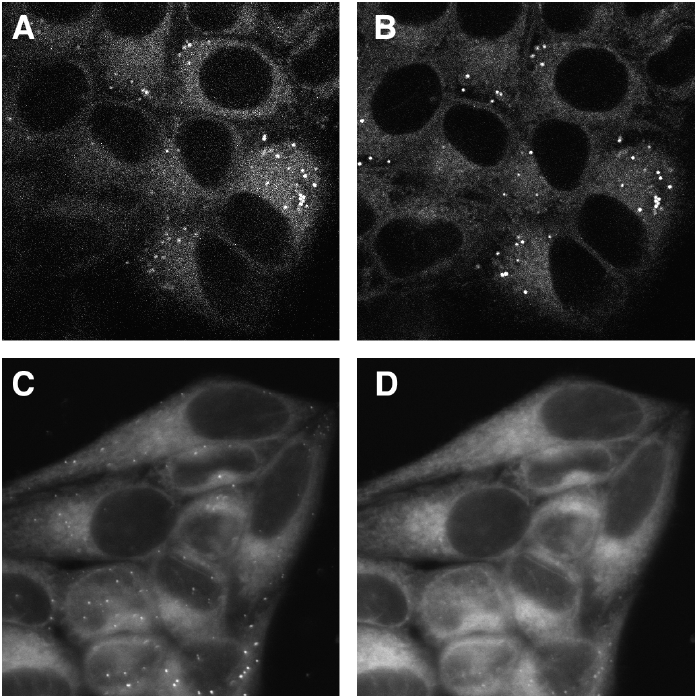
Coumarin fluorescence in living cells after incubation with 200 *µ*M DEACM-DiC8. (A, C) The coumarin fluorescence shows a reticular distribution and is accumulated in punctuate structures. (B) Cells incubated with the lipid droplet marker Nile Red display a similar fluorescence distribution as compared to the coumarin fluorescence shown in panel A. (D) After illumination with UV light, the punctuate structures disappear, whereas the remaining fluorescence maintains a similar pattern as before illumination (compare to panel C). Representative images of at least 3 experiments. Full width of a single image is 74 *µ*m.

### Effect of photoreleased DiC8 on PKCbetaII

It has been shown that classical PKCs can reversibly translocate after stimulating cells either via RTKs or GPCRs. The relocation events are known to coincide with calcium oscillations [5]. Our aim was to understand the role of DAG during these relocation events. Therefore we examined the effect of an instantaneous increase in DAG levels on the relocation of the classical PKC isoform PKCbetaII by using the caged DiC8.

Cells expressing GFP-PKCbetaII and incubated with DEACM-DiC8, were stimulated by a variety of agonists. The reversible relocation events were followed by confocal microscopy and quantified by measuring the fluorescence in the cytoplasm. First, the response of EGFP-PKCbetaII to 100 *µ*M histamine was examined. The results from a representative experiment are shown in figure 5A. The amplitude of the first relocation is relatively high, after which the amplitude of the relocation events decays over time. Notably, equal patterns are observed for calcium oscillations in HeLa cells stimulated with histamine (data not shown). Similar data was obtained from cells incubated with DEACM-DiC8, but not exposed to UV, indicating that the caged compound is not altering the cellular response to histamine (data not shown). However, when cells were briefly exposed to UV after several relocation events, a marked increase in the relocation amplitude was observed, as evident from an increased depletion of PKCbetaII from the cytoplasm (fig. 5B), after which de relocation amplitude decayed again. Similar results were obtained from 7 cells out of a total of 12 cells that displayed oscillations and were exposed to UV.

**Figure 5.**
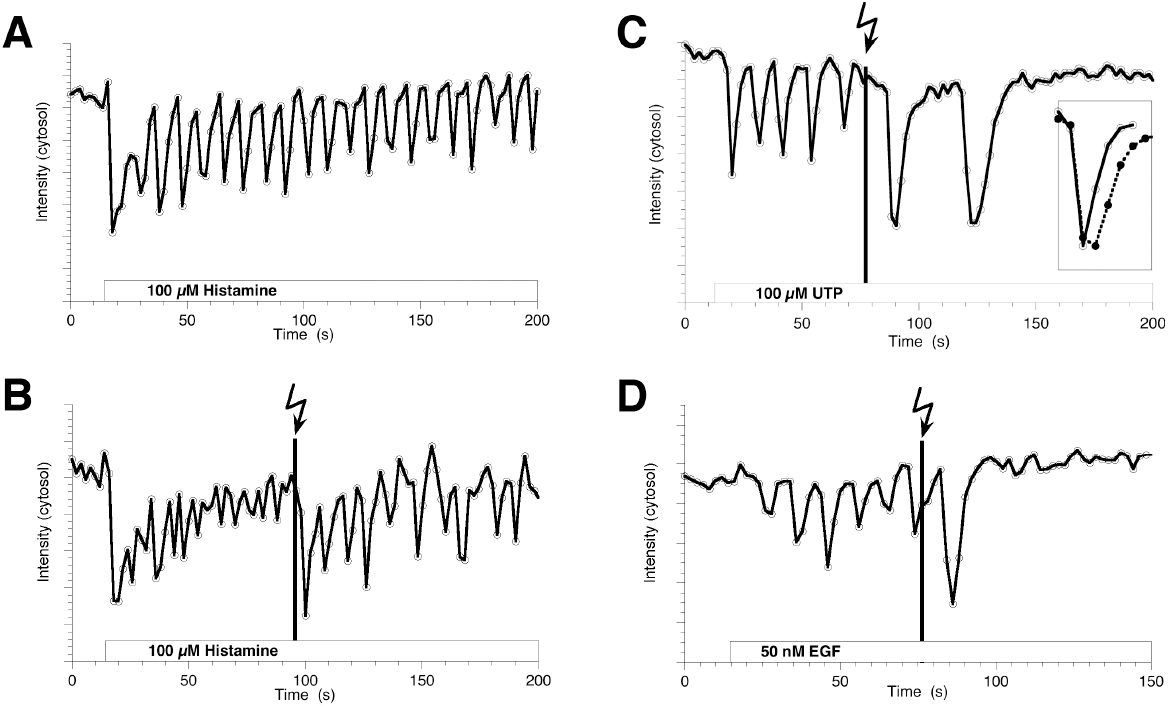
Effect of photoreleased DAG on the reversible relocation of EGFP-PKCbetaII to the plasma membrane induced by receptor activation. The agonists added were 100 *µ*M histamine (A, B), 100 *µ*M UTP (C) and 50 nM EGF (D). The fluorescence intensity of EGFP-PKCbetaII was quantified from the cytoplasm in HeLa cells. After addition of an agonist a clear reduction is observed indicating binding at the plasma membrane. (B-D) The moment of photorelease of DiC8 is indicated by the black line (a single scan with 351/364 nm light, <1s exposure time. Subsequently, the amplitude of the relocation is increased. The inset of panel C compares the amplitude-normalized first relocation event (at t=16s) indicated with open circles with the kinetics of the relocation after uncaging (t=90s) indicated with closed circles.

To examine whether the increased relocation amplitude depends on the agonist that is applied, the same experiments were repeated with the agonist UTP, which stimulates purinergic receptors. As shown in figure 5C, again an increase in relocation amplitude was observed. However, in this case the relocation stopped shortly after UV exposure. The increased relocation amplitude was observed in 13 out 16 cells analyzed. Histamine and UTP both stimulate GPCRs. However, PKCs can also be activated by growth hormones via RTKs. Therefore, experiments were performed in which cells were stimulated with EGF. The results are shown in figure 5D and again an increase in relocation amplitude was observed due to the release of DAG. Similar results were observed in 3 other cells.

## DISCUSSION

The synthesis of a coumarin caged DAG derivative is reported. The coumarin cage provides several advantages due to its intrinsic fluorescence. The caging and uncaging reactions are readily detected on a TLC by UV illumination. Furthermore, the uncaging reaction and the uncaging quantum yield can be determined by measuring the fluorescence increase upon photolysis of DEACM-DiC8. Finally, the compound can be detected in living cells to evaluate loading and localization of the probe.

When applied to living cells, the DEACM-DiC8 is mainly located on endomembranes and accumulates in lipid droplets. Strikingly, the UV-induced photorelease of DAG only increases PKC binding to the plasma membrane during reversible PKC relocation. Since binding of PKC to other membranes was not observed, our results demonstrate that in cells PKC only detects DAG at the plasma membrane. This can be explained by the remarkable preference of PKC for binding negatively charged membranes by interacting with phosphatidylserine [25, 26].

The DAG analog dioctanoylglycerol was selected since it is relatively easy to synthesize and it is more soluble than derivatives with long acyl chains. It should be noted however, that dioctanoylglycerol is not identical to the DAG species that are formed upon PLC activation. Previously, studies have indicated that a variety of DAG species is formed during signaling and that their composition changes over time [27, 28]. Notwithstanding these limitations, we show that the released DiC8 analog is capable of inducing C1a relocation and enhances PKC relocation, clearly demonstrating its biological activity. Still, our method of synthesis should be applicable as well for producing caged derivatives of a variety of DAG analogs with different acyl chain composition. While this was beyond the scope of this study, we believe that such analogs would provide an excellent opportunity to study the DAG specificity for activation of PKCs.

Addition of DiC8 to cells has been used to show that the amplitude of relocation increases for the alpha [29] and gamma [30] PKC isoforms. To the best of our knowledge, this is the first time that a similar effect is observed for the beta isoform. Moreover, our approach demonstrates that the effect of DAG is immediate. Until now it has been difficult to study the kinetics of the response by simply adding DAG to cells, since the response time was mainly determined by the (slow) incorporation of the DAG molecules into the plasma membrane and subsequent flip-flop.

Using the novel caged DAG analog we show that elevated levels of DAG increase the amplitude of PKCbetaII relocation instantaneously, regardless whether cells were stimulated via a receptor tyrosine kinase (EGFR) or via GPCRs (purinergic receptor or histamine H1 receptor). These observations indicate that the effect of DAG on PKC-betaII might be a general mechanism and demonstrate that the levels of DAG produced by PLC during signaling are limiting PKC relocation.

By comparing the kinetics of the relocation events before and after uncaging of DAG, it was observed that besides an increase in the relocation amplitude also the duration of a single relocation event is increased (inset figure 5C). A qualitative inspection of the kinetics of the relocation events reveals that the on-rate is unchanged, whereas the off-rate is decreased by the presence of DAG. Interestingly, previous studies indicated that increasing the DAG concentration delays the dissociation of PKC-gamma [5] and PKC-alpha [29] relative to calcium oscillations.

Since both amplitude and duration of the relocation event are increased, the total number of PKC molecules at the plasma membrane during a calcium oscillation is strongly enhanced. If we assume that the relocation is proportional to kinase activity, the increased DAG levels would effectively amplify the number of phosphorylation events. In this respect it is of note that in some cells the relocation of PKC halted after a few relocation events (e.g. figure 5C and 5D). These results indicate that the elevated levels of DAG subsequently enhance inactivation of the signaling pathway. Interestingly, it was shown that buffering the DAG levels by overexpressing a DAG-binding domain delays the inactivation of PLC [31]. This observation was explained by reduced PKC activation due to lower concentrations of DAG, effectively delaying the desensitation of the signaling pathway. Together, these observations point to a role for DAG in controlling the duration of a signaling event. It is of note that DAG has several effectors besides PKC [32], including TRPC3/TRPC6 [33], protein kinase D [34], chimearin [35, 36] and RasGRP [37]. Therefore, it is not possible to unambiguously identify PKC as the effector that causes inactivation. Future studies will be needed to address the role of other DAG responsive proteins in desensitation of signaling pathways. We feel that the caged DAG analog presented here can be a valuable tool for such studies allowing to evaluate the role of increased levels of DAG at the single cell level in a wide variety of physiological processes. While this work was in progress, the synthesis and application of a number of caged DAG analogues has been reported [38].

## Acknowledgements

We thank Tobias Meyer (Stanford University, Stanford, CA) for providing the plasmid encoding C1a-EGFP and Yusuf Hannun (Duke University Medical Center, Durham, NC) for providing the EGFP-PKCbetaII. This work was supported by a grant (J.G) of the Netherlands Organization for Scientific Research-Council of Chemical Sciences (CW-NWO), a “van der Leeuw” grant (T.W.J.G.) of Netherlands Organization for Scientific Research (NWO) and the EU integrated project on “Molecular Imaging” LSHG-CT-2003-503259.

## REFERENCES

[1] S.G. Rhee, Regulation of phosphoinositide-specific phospholipase C, Annu. Rev. Biochem. 70 (2001) 281–312.

[2] M.J. Berridge, R.F. Irvine, Inositol trisphosphate, a novel second messenger in cellular signal transduction, Nature 312 (1984) 315–321.

[3] A.C. Newton, Protein kinase C: structural and spatial regulation by phosphorylation, cofactors, and macromolecular interactions, Chem. Rev. 101 (2001) 2353–2364.

[4] X. Feng, J. Zhang, L.S. Barak, T. Meyer, M.G. Caron, Y.A. Hannun, Visualization of dynamic trafficking of a protein kinase C betaII/green fluorescent protein conjugate reveals differences in G protein-coupled receptor activation and desensitization, J Biol Chem 273 (1998) 10755–10762.

[5] E. Oancea, T. Meyer, Protein kinase C as a molecular machine for decoding calcium and diacylglycerol signals, Cell 95 (1998) 307–318.

[6] N. Sakai, K. Sasaki, N. Ikegaki, Y. Shirai, Y. Ono, N. Saito, Direct visualization of the relocation of the gamma-subspecies of protein kinase C in living cells using fusion proteins with green fluorescent protein, J. Cell Biol. 139 (1997) 1465–1476.

[7] S.R. Adams, R.Y. Tsien, Controlling cell chemistry with caged compounds, Annu. Rev. Physiol. 55 (1993) 755–784.

[8] J.C. Lenz, H.P. Reusch, N. Albrecht, G. Schultz, M. Schaefer, Ca2+-controlled competitive diacylglycerol binding of protein kinase C isoenzymes in living cells, J. Cell Biol. 159 (2002) 291–302.

[9] G. Reither, M. Schaefer, P. Lipp, PKCalpha: a versatile key for decoding the cellular calcium toolkit, J. Cell Biol. 174 (2006) 521–533.

[10] J.Q. He, Y. Pi, J.W. Walker, T.J. Kamp, Endothelin-1 and photoreleased diacylglycerol increase L-type Ca2+ current by activation of protein kinase C in rat ventricular myocytes, The Journal of physiology 524 Pt 3 (2000) 807–820.

[11] X.P. Huang, R. Sreekumar, J.R. Patel, J.W. Walker, Response of cardiac myocytes to a ramp increase of diacylglycerol generated by photolysis of a novel caged diacylglycerol, Biophys. J. 70 (1996) 2448–2457.

[12] V.G. Robu, E.S. Pfeiffer, S.L. Robia, R.C. Balijepalli, Y. Pi, T.J. Kamp, J.W. Walker, Localization of functional endothelin receptor signaling complexes in cardiac transverse tubules, J Biol Chem 278 (2003) 48154–48161.

[13] T. Furuta, M. Iwamura, New caged groups: 7-substituted coumarinylmethyl phosphate esters, Methods Enzymol. 291 (1998) 50–63.

[14] V. Hagen, S. Frings, J. Bendig, D. Lorenz, B. Wiesner, U.B. Kaupp, Fluorescence spectroscopic quantification of the release of cyclic nucleotides from photocleavable [bis(carboxymethoxy)coumarin-4-yl]methyl esters inside cells, Angew. Chem. Int. Ed. Engl. 41 (2002) 3625–3628, 3516.

[15] R.O. Schonleber, J. Bendig, V. Hagen, B. Giese, Rapid photolytic release of cytidine 5'-diphosphate from a coumarin derivative: a new tool for the investigation of ribonucleotide reductases, Bioorg. Med. Chem. 10 (2002) 97–101.

[16] T. Furuta, S.S. Wang, J.L. Dantzker, T.M. Dore, W.J. Bybee, E.M. Callaway, W. Denk, R.Y. Tsien, Brominated 7-hydroxycoumarin-4-ylmethyls: photolabile protecting groups with biologically useful cross-sections for two photon photolysis, Proc. Natl. Acad. Sci. U.S.A. 96 (1999) 1193–1200.

[17] R. Sreekumar, Y.Q. Pi, X.P. Huang, J.W. Walker, Stereospecific protein kinase C activation by photolabile diglycerides, Bioorg. Med. Chem. Lett. 7 (1997) 341–346.

[18] J. Goedhart, T.W.J. Gadella, Jr., Photolysis of caged phosphatidic acid induces flagellar excision in Chlamydomonas, Biochemistry 43 (2004) 4263–4271.

[19] A.Z. Suzuki, T. Watanabe, M. Kawamoto, K. Nishiyama, H. Yamashita, M. Ishii, M. Iwamura, T. Furuta, Coumarin-4-ylmethoxycarbonyls as phototriggers for alcohols and phenols, Org Lett 5 (2003) 4867–4870.

[20] V. Hagen, J. Bendig, S. Frings, T. Eckardt, S. Helm, D. Reuter, U.B. Kaupp, Highly Efficient and Ultrafast Phototriggers for cAMP and cGMP by Using Long-Wavelength UV/Vis-Activation, Angew. Chem. Int. Ed. Engl. 40 (2001) 1045–1048.

[21] B. Schade, V. Hagen, R. Schmidt, R. Herbrich, E. Krause, T. Eckardt, J. Bendig, Deactivation Behavior and Excited-State Properties of (Coumarin-4-yl)methyl Derivatives. 1. Photocleavage of (7-Methoxycoumarin-4-yl)methyl-Caged Acids with Fluorescence Enhancement, J. Org. Chem. 64 (1999) 9109–9117.

[22] T. Eckardt, V. Hagen, B. Schade, R. Schmidt, C. Schweitzer, J. Bendig, Deactivation Behavior and Excited-State Properties of (Coumarin-4-yl)methyl Derivatives. 2. Photocleavage of Selected (Coumarin-4-yl)methyl-Caged Adenosine Cyclic 3',5' Monophosphates with Fluorescence Enhancement, J. Org. Chem. 67 (2002) 703–710.

[23] E. Oancea, M.N. Teruel, A.F. Quest, T. Meyer, Green fluorescent protein (GFP)-tagged cysteine-rich domains from protein kinase C as fluorescent indicators for diacylglycerol signaling in living cells, J. Cell Biol. 140 (1998) 485–498.

[24] P. Greenspan, E.P. Mayer, S.D. Fowler, Nile red: a selective fluorescent stain for intracellular lipid droplets, J. Cell Biol. 100 (1985) 965–973.

[25] J.E. Johnson, J. Giorgione, A.C. Newton, The C1 and C2 domains of protein kinase C are independent membrane targeting modules, with specificity for phosphatidylserine conferred by the C1 domain, Biochemistry 39 (2000) 11360–11369.

[26] M. Medkova, W. Cho, Differential membrane-binding and activation mechanisms of protein kinase C-alpha and -epsilon, Biochemistry 37 (1998) 4892–4900.

[27] S. Carrasco, I. Merida, Diacylglycerol, when simplicity becomes complex, Trends Biochem. Sci. 32 (2007) 27–36.

[28] M.N. Hodgkin, T.R. Pettitt, A. Martin, R.H. Michell, A.J. Pemberton, M.J. Wakelam, Diacylglycerols and phosphatidates: which molecular species are intracellular messengers?, Trends Biochem. Sci. 23 (1998) 200–204.

[29] A. Tanimura, A. Nezu, T. Morita, N. Hashimoto, Y. Tojyo, Interplay between calcium, diacylglycerol, and phosphorylation in the spatial and temporal regulation of PKCalpha-GFP, J Biol Chem 277 (2002) 29054–29062.

[30] G. Halet, R. Tunwell, S.J. Parkinson, J. Carroll, Conventional PKCs regulate the temporal pattern of Ca2+ oscillations at fertilization in mouse eggs, J. Cell Biol. 164 (2004) 1033–1044.

[31] F. Codazzi, M.N. Teruel, T. Meyer, Control of astrocyte Ca(2+) oscillations and waves by oscillating relocation and activation of protein kinase C, Curr. Biol. 11 (2001) 1089–1097.

[32] N. Brose, C. Rosenmund, Move over protein kinase C, you've got company: alternative cellular effectors of diacylglycerol and phorbol esters, J. Cell Sci. 115 (2002) 4399–4411.

[33] T. Hofmann, A.G. Obukhov, M. Schaefer, C. Harteneck, T. Gudermann, G. Schultz, Direct activation of human TRPC6 and TRPC3 channels by diacylglycerol, Nature 397 (1999) 259–263.

[34] C.L. Baron, V. Malhotra, Role of diacylglycerol in PKD recruitment to the TGN and protein transport to the plasma membrane, Science 295 (2002) 325–328.

[35] S. Ahmed, R. Kozma, C. Monfries, C. Hall, H.H. Lim, P. Smith, L. Lim, Human brain n-chimaerin cDNA encodes a novel phorbol ester receptor, Biochem. J. 272 (1990) 767–773.

[36] M.J. Caloca, N. Fernandez, N.E. Lewin, D. Ching, R. Modali, P.M. Blumberg, M.G. Kazanietz, Beta2-chimaerin is a high affinity receptor for the phorbol ester tumor promoters, J Biol Chem 272 (1997) 26488–26496.

[37] J.O. Ebinu, D.A. Bottorff, E.Y. Chan, S.L. Stang, R.J. Dunn, J.C. Stone, RasGRP, a Ras guanyl nucleotide- releasing protein with calcium- and diacylglycerol-binding motifs, Science 280 (1998) 1082–1086.

[38] A. Nadler, G. Reither, S. Feng, F. Stein, S. Reither, R. Müller, C. Schultz, The fatty acid composition of diacylglycerols determines local signaling patterns, Angew. Chem. Int. Ed. Engl. 10 (2013) 6330–6334

